# Clinical and Genetic Determinants of Glioblastoma Multiforme Survival: A Retrospective Analysis

**DOI:** 10.1101/2025.01.23.634459

**Authors:** Julia L. Gutiérrez-Arroyo, Pia Gallego-Porcar, Elvira Carbonell-Martinez, Luis G. González-Bonet, Maria Victoria Ibañez, María Díaz-Ruiz, Hugo Caballero-Arzapalo, Ariadna Soto, Guillermo Garcia-Oriola, Jose Maria Borras-Moreno, Conrado Martinez-Cadenas, Maria Angeles Marques-Torrejon

## Abstract

Glioblastoma, the most aggressive primary brain tumour in adults, has a poor prognosis and limited survival despite advances in treatment. This study analysed 61 patients with glioblastoma multiforme treated at the General University Hospital of Castellon, Spain, focusing on clinical, tumour-specific and genetic factors influencing disease outcome. Variables included age, sex, BMI, extent of surgical resection, and use of radiotherapy or chemotherapy. Tumour characteristics assessed included location, size, proximity to the ventricular system and surgical approach. Genetic mutations in the *IDH, EGFR, TP53* and *CDKN2A* genes were also analysed. Kaplan-Meier analysis was used to assess the impact of these factors on overall survival and progression-free survival. A significant finding was the strong association between surgical approach, tumour proximity to the ventricular system and survival: patients with tumours closer to the ventricles had significantly shorter survival, highlighting the critical role of spatial tumour characteristics in glioblastoma multiforme outcomes. These results suggest that integrating clinical, genetic and spatial tumour data into personalised treatment approaches could improve prognosis. Understanding these factors is critical to developing more effective strategies to meet the challenges of this aggressive and complex disease.

## INTRODUCTION

Glioblastoma (GB), classified as a grade 4 astrocytoma by the World Health Organization (WHO)^1,2^, is the most prevalent, aggressive, and lethal primary brain tumour in adults. GB accounts for 50.1% of all malignant brain tumours, with an annual global incidence of approximately 3 cases per 100,000 individuals. The median age at diagnosis is around 64 years, with incidence rates increasing with age and peaking at 15 cases per 100,000 individuals between 75 and 84 years old^3,4^. GB is 1.6 times more common in males than females and occurs more frequently in Caucasians compared to other ethnic groups^4^. Known for its rapid growth, high invasiveness, and significant molecular diversity^2^, GB typically arises in the cerebral hemispheres. About 95% of cases occur in the supratentorial region, particularly within the frontal and temporal lobes^5^. Despite advances in treatment, including extensive surgical resection, radiotherapy, and temozolomide-based chemotherapy (Stupp protocol)^6^, the median survival for GB patients remains dismal at 12–15 months post-diagnosis. This underscores the urgent need for novel therapies that effectively target the biological and molecular complexity of GB^7,8^.

GB has extensive tumour heterogeneity and plasticity at the cytopathological, transcriptional, and genetic levels^9-12^. Moreover, its high permeability and blood-brain barrier (BBB) protection pose significant therapeutic challenges^8,13,14^. Glioma stem cells (GSCs) are a rare population within GBs characterized by genetic instability, self-renewal capacity, tumour-initiating potential, and the ability to differentiate into diverse cellular subsets, contributing to the heterogeneity of GB^15-20^. Moreover, GSCs are resistant to apoptosis^21-24^, can influence various components of the tumour microenvironment, play a role in activating angiogenesis and immunosuppression, and contribute to radio- and chemo-resistance. Additionally, this resistance can be induced by hypoxic conditions within the tumour microenvironment, which promote the maintenance of stem-like properties in GSCs, further enhancing their ability to evade treatments and support tumour progression^25,26^. The fact that current treatments cannot completely eliminate certain subsets of GSCs is considered a major reason why the tumour almost always comes back after therapy^27^.

Recent advances in genomic profiling have provided deeper insights into the genetic alterations driving GB. *IDH1*/*2* mutations are predominantly found in grade IV astrocytoma, formerly known as secondary GBs, and lower-grade gliomas^28^. Secondary GBs are different from primary GBs because they develop from pre-existing lower-grade gliomas (classified as WHO grades 2 or 3) instead of appearing out of nowhere, like primary GBs do^29^. Lower-grade gliomas tend to grow more slowly and have better survival rates initially, but over time, they can pick up additional genetic changes, such as mutations in *TP53* or loss of *ATRX*, which can eventually lead to their progression into secondary GBs^30^ This distinction between primary and secondary GBs is crucial because it highlights the differences in their molecular and clinical pathways^31^. In contrast, *EGFR* amplification, present in approximately 40% of GB cases, is linked to worse outcomes by promoting aggressive tumour behaviour through constitutive activation of growth signalling pathways^32^. *TP53* mutations, affecting the p53 pathway in 87% of cases, significantly contribute to disease progression, while alterations in *CDKN2A*, a key cell cycle regulator gene, further impair apoptosis and disrupt cell cycle control^33^. *RTK/PI3K/PTEN* pathway alterations are observed in 88% of GB cases, emphasizing their role in tumorigenesis. Additionally, loss of heterozygosity on chromosome 10 is one of the most common chromosomal abnormalities identified^34^. These genetic insights not only enhance our understanding of GB biology but also have potential to improve diagnosis, predict outcomes, and inform personalized therapies. For example, *IDH* mutations offer prognostic value, while *EGFR, TP53*, and *CDKN2A* alterations highlight the molecular complexity and therapeutic resistance of GB^35^.

The development of GB is influenced not only by genetic factors but also by clinical and lifestyle variables, including patient sex, age, and overall health status. Studies suggest that sex may affect survival, with some evidence indicating slightly longer survival times in female patients^36^. Lifestyle factors such as smoking, family cancer history, and viral infections (e.g., hepatitis B, COVID-19) may also influence disease progression and treatment response, although their roles in GB remain underexplored^37^.

Tumour location relative to brain ventricles has emerged as a critical prognostic factor, though findings on its impact on overall survival (OS) have been inconsistent^38^. While GB distance from the subventricular neural stem cell niche has not been correlated with survival^39^, a recent meta-analysis reported that GBs in contact with the lateral ventricle are associated with lower survival. This effect may be independent of established survival predictors, emphasizing the clinical relevance of the ventricular-subventricular zone contact in GB biology^37,39^. Furthermore, tumour location near the third ventricle and the contrast-enhancing tumour border has been identified as a prognostic factor, particularly in elderly patients^40^. Understanding these spatial relationships and their biological implications is crucial for devising more effective therapies and improving GB prognosis.

The aim of this study was to explore the different factors that might affect survival in patients with GB. We wanted to study factors such as demographic characteristics, clinical and surgical factors, and even genetic aspects to see which ones really make a difference in survival. We also wanted to understand how tumour characteristics, treatments, and recurrence patterns play a role in survival. Through this analysis, our goal was to enhance the understanding of factors that could inform prognosis and guide the development of more effective treatment strategies for GB.

## RESULTS

### Demographic and Clinical Variables

In this study, we looked at 61 patients who were diagnosed with GB. The median age at the time of diagnosis was 61.5 years, with ages ranging from 16 to 80 years. Interestingly, there was an almost even split between male and female patients, with a 1:1 ratio. The average BMI (Body Mass Index) of the group was 26.6 kg/m^2^, with a standard deviation of ±4.2. When it came to survival, the median OS was 13.7 months, ranging from as little as 3 months to as long as 32 months. The median progression-free survival (PFS) was slightly longer, at 12.4 months, with a range of 1 to 18 months.

Using Kaplan-Meier survival curve, we also analysed whether demographic factors like sex and age (Figure 1A), or BMI had any impact on survival. From this analysis, we found no significant differences in OS or PFS between male and female patients (Figure 1B). Similarly, no clear link was found between survival outcomes and other factors like age (Figure 1A) or BMI (data not shown).

**Figure 1.**
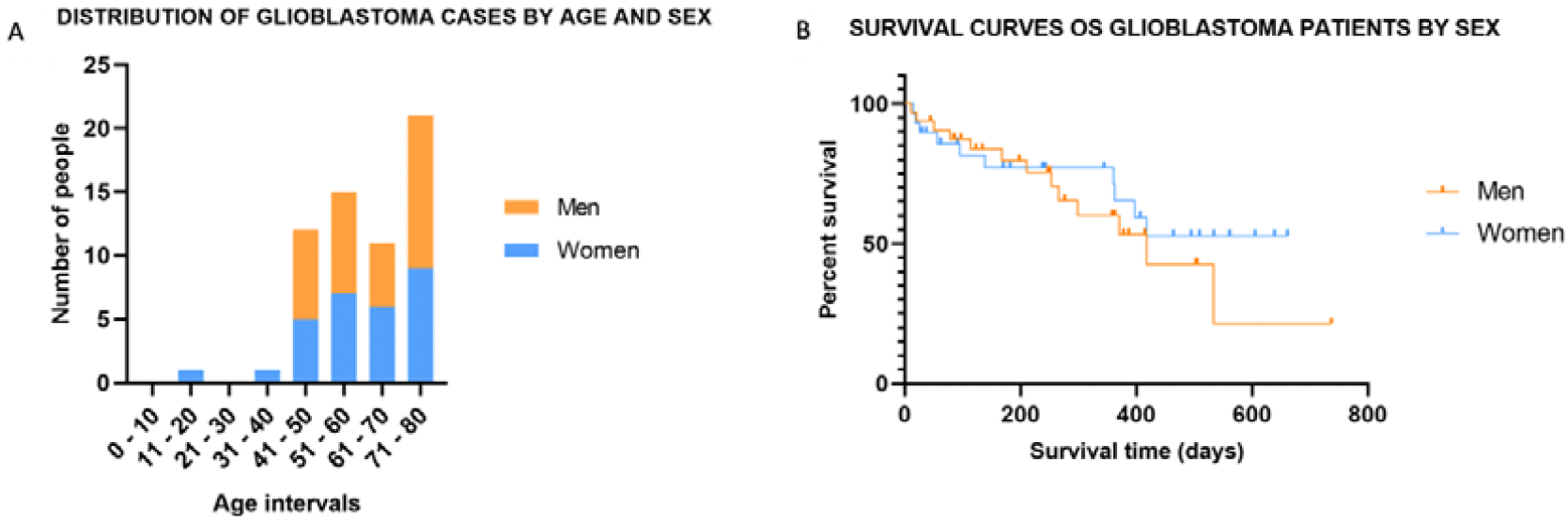
Distribution and survival of GB cases by age and sex. (A) The histogram shows the number of cases across age intervals (in years) divided by sex: men (orange) and women (blue). The highest incidence is observed in the 61–70 and 71–80 age groups. (B) OS curves for glioblastoma patients stratified by sex. Kaplan-Meier curves represent survival times for men (orange line) and women (blue line).

### Tumour Features and Treatment Details

Tumour locations in this study varied, but most were found in the frontal and temporal lobes. When we looked at treatment outcomes, surgical resection stood out as a key factor influencing OS. More than 50% of patients who underwent gross total resection were still alive at the end of the follow-up period; therefore, this group did not reach the median OS during the study period (n=16). In contrast, the OS for patients who underwent subtotal resection was 13.7 months (n=25), and for those who only underwent a biopsy, it was 8.72 months (n=19), with the difference being statistically significant (p < 0.043) (Figure 2C). This highlights how important it is to remove as much of the tumour as possible during surgery. That said, it is also important to keep in mind that the extent of resection often depends on where the tumour is located, since some areas of the brain are harder to access safely.

**Figure 2.**
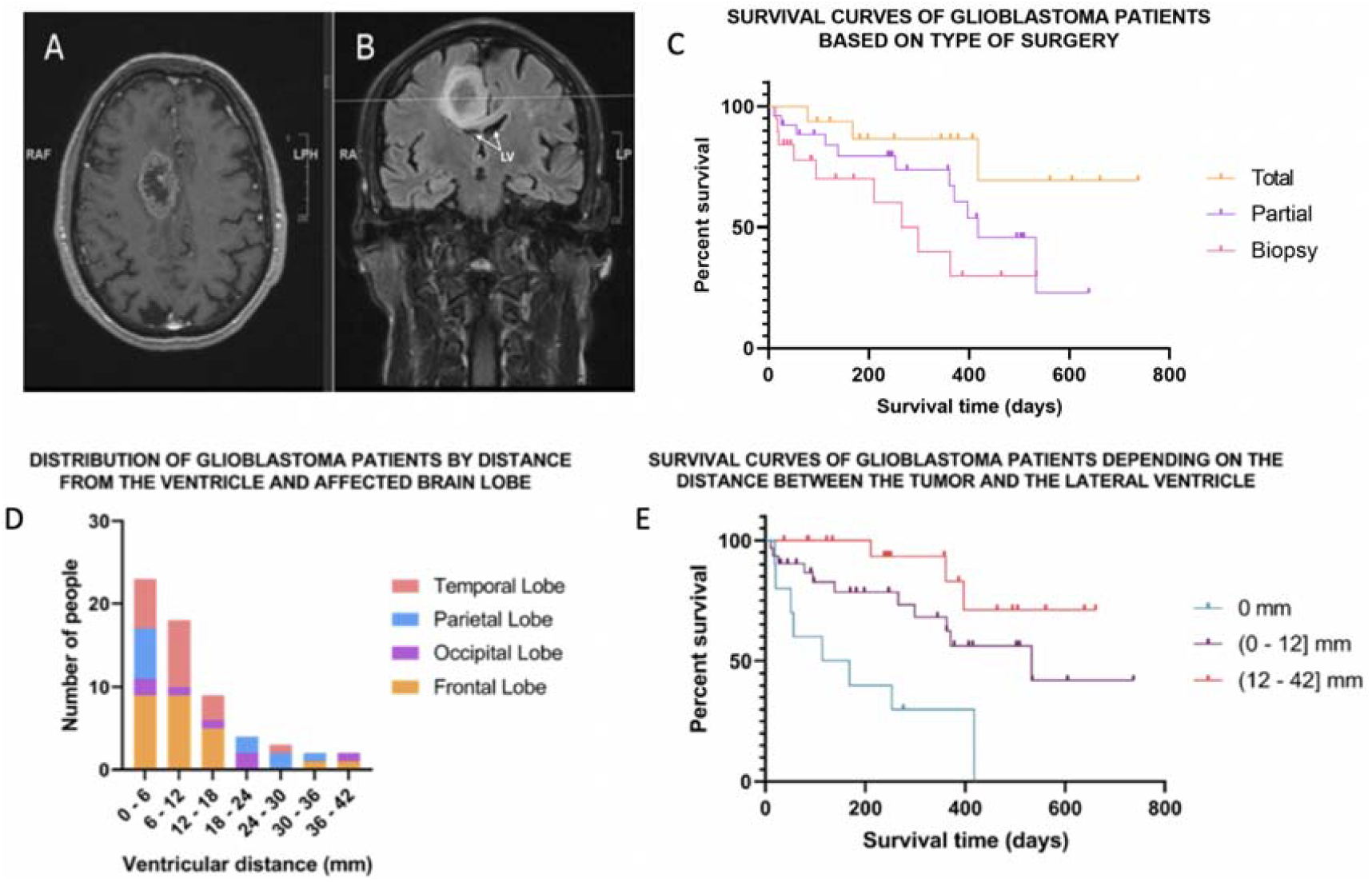
(A, B) MRI (Magnetic resonance imaging) scans showing the GB location relative to the lateral ventricle in axial (A) and coronal (B) views, highlighting the tumour’s proximity to ventricular structures. (C) OS curves of GB patients stratified by type of surgery: total resection, partial resection, and biopsy. The orange line represents total resection, the purple line represents partial resection, and the red line represents biopsy. (D) Distribution of GB cases by ventricular distance (mm) and affected brain lobe (temporal, parietal, occipital, and frontal). (E) Kaplan-Meier survival curves of GB patients stratified by tumour distance to the lateral ventricle: 0 mm (direct contact), 0–12 mm, and 12–42 mm. Survival is significantly reduced for tumours that are closer to the ventricle.

Interestingly, treatments like radiotherapy (n=56) and chemotherapy (n=59), including the widely used Stupp protocol, did not show significant differences in OS in this particular group of patients (data not shown). This suggests that variations in survival were more closely tied to surgical outcomes and specific tumour characteristics rather than differences in these additional treatments.

### Effect of recurrence, tumour location and distance of the tumour to the ventricle on OS

Out of all 61 patients included in the study, tumour recurrence was observed in 22 cases (36.1%) during the study period. The median time to recurrence was relatively short, at 3.4 months. Secondary surgeries were performed in 12 patients but, unfortunately, these additional procedures did not have a significant impact on survival outcomes (data not shown).

When tumour location (Figure 2D) in the brain was analysed in relation to OS, no meaningful correlations were found. This suggests that survival outcomes are not strongly influenced by the specific lobe or region of the brain where the tumour was located (p = 0.220) (Figure 2D).

The proximity of the tumour to the ventricular system (Figure 2A and 2B) also showed a strong and significant association with survival (p < 0.001). For Kaplan-Meier survival analysis, patients were also divided into three groups based on the distance between the tumour and the ventricular system: tumours directly in contact with the ventricle (0 mm), tumours located 0–12 mm away, and tumours 12–42 mm away. Patients whose tumours were in direct contact with the ventricle had the shortest median OS (3.74 months), followed by those with tumours located 0–12 mm away (17.48 months). Interestingly, 50% of patients with tumours situated 12–42 mm away from the ventricle were still alive at the end of the follow-up period (therefore, median OS was not reached for this group during the study period) (Figure 2E).

### Genetic Analysis

In this study, we did not find any significant links between survival outcomes and mutations in the analysed genes —*IDH1, EGFR, TP53*, and *CDKN2A*. Likewise, copy number variations (CNVs) in *EGFR, CDKN2A, FGFR2*, and *PTEN* did not show any significant differences in OS or PFS. Even when looking at RNA expression levels or the MGMT (O6-methylguanine–DNA methyltransferase) promoter methylation status, there were no significant associations with survival (data not shown).

We also performed Kaplan-Meier survival analysis to look at how individual mutations might affect OS and PFS, but none of the genetic alterations we examined showed a significant impact (Figure 4A-D). In addition, we analysed whether combinations or interactions of the different mutations, for example co-occurring (data not shown) or mutually exclusive mutations in *IDH1, EGFR, TP53*, and *CDKN2A* had any influence on survival outcomes. Unfortunately, these analyses also did not reveal any meaningful associations (data not shown).

**Figure 4.**
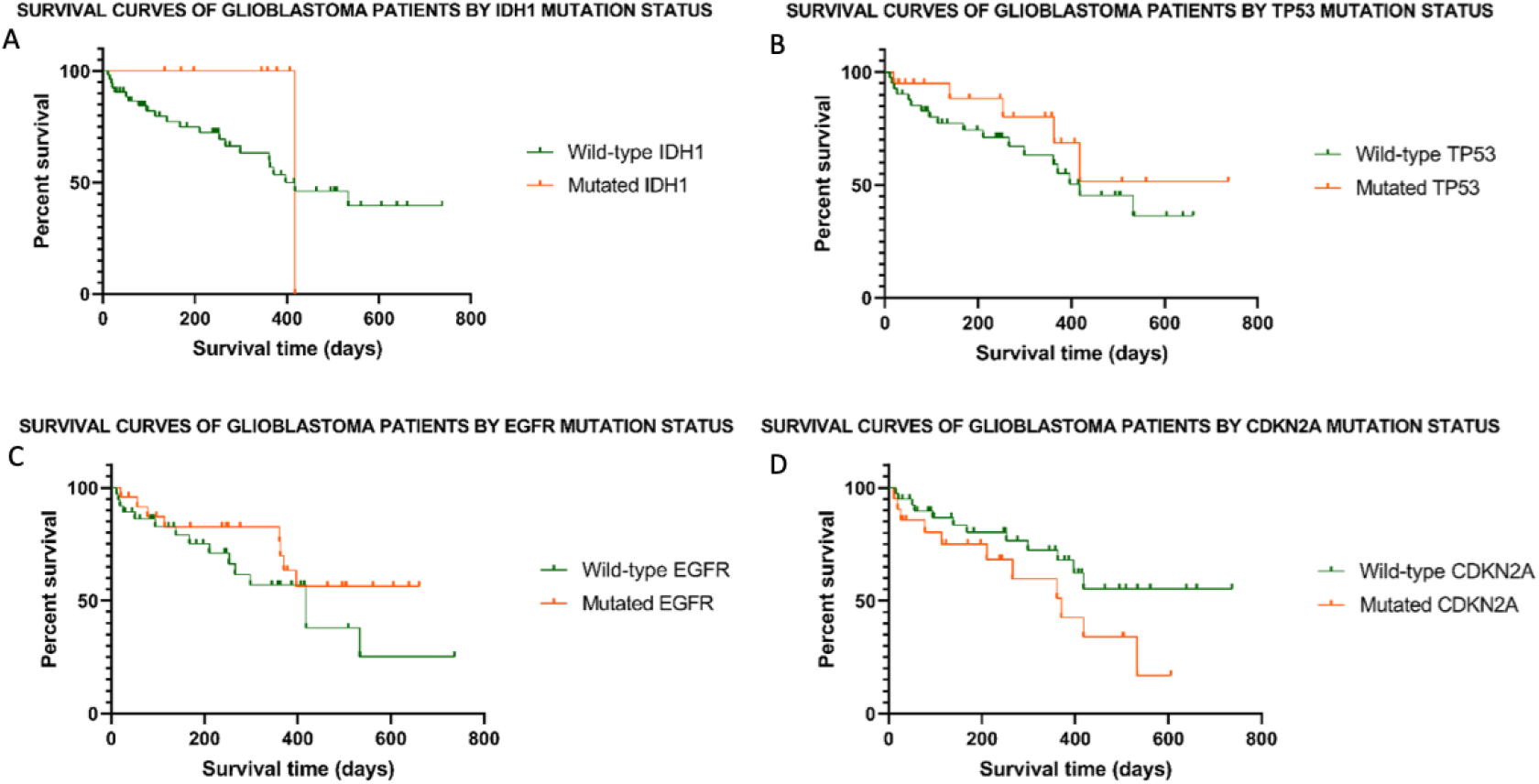
Kaplan-Meier survival curves for GB patients stratified by mutation status of key genes: (A) *IDH1*, (B) *TP53*, (C) *EGFR*, and (D) *CDKN2A*. Wild-type and mutated forms shown in green and orange, respectively.

These results suggest that, at least within this group of patients, the genetic changes we assessed do not provide any useful insights into survival outcomes.

### Lifestyle and Medical History

Lifestyle and medical history factors, such as smoking habits, previous cancer diagnoses, a history of COVID-19 infection, hepatitis B status, or Type II diabetes, did not show any significant impact on survival outcomes in this group of patients (data not shown).

## DISCUSSION

In this study, we explored various factors that could influence survival in patients with glioblastoma (GB), including various patient-related, tumour-related, and treatment-related factors. Most of the factors analysed did not show an effect on patient median overall survival (OS) or median progression-free survival (PFS), perhaps due to the small number of patients. However, the distance of the tumour to the ventricles, as well as surgery type had a strong correlation with survival. The proximity of the tumour to the lateral ventricle, as well as the observed difference in survival rates, lead to the hypothesis that the proximity of the tumour to the cerebrospinal fluid (CSF) could be a contributing factor to the progression of the disease. The corpus callosum, a myelin structure that connects the two lateral hemispheres of the brain, is also attached to the lateral ventricle and could act as a conduit for cancer cell invasion. Furthermore, this hypothesis could suggest that a reservoir of neural stem cells is present within the lateral ventricle of the brain, which could act as a reservoir for cancer cells within the tumour. This fact could be assessed by the glioma stem cell marker CD133^41^.

When we examined demographic factors such as age, sex, and BMI, we found that none of them had a significant impact on survival outcomes. This is largely in line with other studies^42^, which suggest that these factors are less critical for predicting overall survival in GB compared to other variables, such as treatment options or tumour behaviour^43^. As for treatments like radiotherapy and chemotherapy, including the Stupp protocol, they did not seem to significantly impact survival. This could be due to GB complexity, since different patients respond to treatment in very different ways, making it harder to draw clear conclusions. Regarding tumour location, most were in the frontal and temporal lobes. However, surprisingly, specific tumour location did not seem to affect patient survival.

Regarding genetic factors, mutations in GB driver genes such as *IDH1, EGFR, TP53*, and *CDKN2A*, or CNVs in genes like *EGFR* and *PTEN*, did not show any strong connection to survival outcomes in this study. It is likely that the absence of a significant difference in *IDH* mutation is attributable to the limited size of the mutated group. Furthermore, it should be noted that the current classification of GB does not include *IDH* mutation as a diagnostic criterion. Even other factors like MGMT promoter methylation and RNA expression levels did not seem to give us any useful information about survival. While other studies^31-33^ have found links between certain genetic markers and survival in GB, we did not observe the same results. This could be due to patient differences or because our sample size was too small to pick up on subtle effects, but it is clear that we still lack considerable insight on how genetic changes impact GB.

Recurrence was also highly relevant. About 76% of patients had tumour relapses, with an average time to recurrence of around 6 months. Unfortunately, secondary surgeries for recurrent tumours did not seem to improve survival, which really highlights how limited our treatment options are when GB reappears. We need better strategies for dealing with recurrent GB, something that is clearly still a massive challenge in the field.

On the other hand, when we looked at lifestyle and medical history factors like smoking, past cancer, COVID-19 infection, and conditions like hepatitis B and Type II diabetes, they did not seem to affect survival at all. This suggests that these factors might not matter as much in the context of GB. However, in GB patients who are motivated to make lifestyle adjustments to improve their outcomes, the exciting news is that dietary restriction of sugar and caloric intake in particular seems to show some promise, with even more good news coming from exercise, vitamin supplementation, and cannabis use showing potential benefits as well^37,44^.

However, the most important factor in patient survival was how much of the tumour was removed during surgery. Patients who had a gross total resection, in which as much of the tumour as possible is removed, lived much longer (around 18 months) compared to those who only had partial resection (12 months) or just a biopsy (9 months). This highlights the value surgery has on GB survival time, though it is important to remember that tumour location can make it harder to remove the totality of tumour tissue, especially if it is located near critical structures^45,46^. A significant challenge in the management of glioblastoma (GB) is the difficulty neurosurgeons face in accessing the tumour. The primary objective is to achieve complete removal of the tumour without compromising the patient’s life. However, this is not without its challenges, as it is not possible to thoroughly cleanse adjacent areas due to the potential compromise to life^47^. This is a consequence of the high invasiveness of brain cells, which renders total surgery a viable option for increasing survival but not preventing relapse.

Perhaps the most relevant finding of this work is that tumours located close to the ventricular system are associated with shorter survival. Patients with tumours that were in direct contact with the ventricles had the shortest survival times, which suggests that where the tumour is located in relation to the ventricles might influence tumour growth or treatment efficacy. There is a growing interest in assessing the effectiveness of VSVZ radiation in addition to the standard of care treatment for GB. For instance, one randomised trial is currently underway (ClinicalTrials.gov Identifier: NCT02177578), and our results may inform the analysis of such trials. For example, in current trials, it may be beneficial to test the differential effectiveness of VSVZ radiation in niche-contacting and non-contacting GBs.

In conclusion, the findings of this study really highlight the importance of surgery in treating GB, especially when it comes to how much of the tumour can be safely removed. While genetic factors did not provide us with clear answers, it is still important to keep studying them, as they could hold the key to better treatments in the future. The relationship between tumour location and survival is also something that deserves more attention, as it could help us improve prognostic models. Overall, there is still a lot we need to figure out, but the hope is that by continuing to explore new treatment options and gathering more data, we can find better ways to help patients with GB in the future.

## METHODS

### Study Design and Patient Selection

This retrospective cohort study includes 61 adult patients diagnosed with GB at the Castellon University General Hospital, Castellon, Spain. Patients were selected based on confirmed GB diagnosis and the availability of complete clinical and genetic data. All results reflect data up to February 12, 2023, with patient survival information updated to this date.

### Data Collection

For each patient, we created a database including different factors:

1. Demographic and Clinical Variables: sex, age at diagnosis, year of birth, BMI (calculated from weight and height), and survival metrics (OS, PFS).
2. Tumour and Treatment Details: tumour location (frontal, parietal, temporal, occipital), tumour subtype, and treatment information including radiotherapy, extent of surgical resection (total, partial or subtotal, biopsy), and chemotherapy regimen (Stupp protocol with temozolomide, PCV regimen, and adjunctive use of levetiracetam/Bevacizumab). In addition, the distance of the tumour to the ventricular system was measured using image analysis within the software associated with the fMRI scans of each patient. The image with the highest tumour density was selected for measurement. For statistical analysis, distances were categorized into three groups: tumours directly contacting the ventricle (0 mm), tumours located near the ventricle (0– 12 mm), and tumours located farther from the ventricle (12–42 mm).
3. Recurrence Data: The recurrence type, date, time to recurrence, and details of any secondary surgical intervention were collected. Additionally, the distance from the tumour to the nearest ventricle.
4. Lifestyle and Medical History: Presence of prior cancer, smoking status, history of COVID-19 infection, hepatitis B status, and Type II diabetes.
5. Genetic Analysis: NGS was performed in tumour DNA and tumour RNA on a panel of target genes. DNA and RNA extraction was performed automatically using QiAcube extractor (Qiagen, Hilden, Germany) following the manufacturer’s instructions. Massive NGS sequencing was performed with Ion Torrent technology (Ion Torrent™ Genexus™ Integrated Sequencer) from Thermo Fisher Scientific (Waltham, MA, USA). The Oncomine Precision panel - GX5 - Solid Tumour - w3.2.0 DNA and Fusions Panel was used with target regions defined in Target Regions Oncomine precision v3.6.20210407.designed.bed. The genetic sequencing data was analysed at the Department of Clinical Analysis of the Castellon General University Hospital, following their standard protocol for post-surgery tumour examination. In our case, given the diagnosis of GB, the analysis focused on identifying mutations commonly associated with this type of tumour.

The bioinformatics platform used was Genexus System. The detected variants have been filtered and visualized with the Ion Reporter™ Software 5.18 and IGV Integrative Genomics Viewer programs. The detection limit of the technique is VFA>0.5%. The cancer panel sequenced included the following genes, where two types of genetic mutations were determined: i) cancer driving mutations in the following genes: *IDH1, IDH2, EGFR, TP53, KRAS, HRAS, RET, PTEN, NTRK1, PIK3CA, MAP2K1*, and *BRAF*; and ii) copy number variations (CNVs) in the following genes *EGFR, CDKN2A, FGFR2, PTEN, AR*. Additionally, RNA expression levels and MGMT promoter methylation status were also performed.

### Statistical Analysis

SPSS software was used for all statistical analyses.

Kaplan-Meier survival analysis was performed to assess OS and PFS in relation to the following factors: i) clinical and treatment variables — sex, radiotherapy, chemotherapy, surgical type (total, partial, biopsy), tumour location (frontal, parietal, temporal, occipital), distance from the tumour to the ventricular system, and tumour size —; ii) genetic factors — presence of somatic mutations in *IDH, EGFR, TP53*, and *CDKN2A* —; and iii) interaction analysis — pairwise interactions between key genetic mutations (*IDH, EGFR, TP53, CDKN2A*) were also evaluated for potential synergistic or antagonistic effects on survival outcomes.

Survival curves were compared across subgroups using the log-rank test, and multivariate Cox regression models were employed to identify factors independently associated with survival. Statistical significance was set at p < 0.05.

## AUTHOR CONTRIBUTIONS STATEMENT

LGG-B, CM-C and MAM-T conceived the study; LGG-B and MAM-T planned and designed the research; MD-R performed the genetic analyses; JLG-A, PG-P and EC-M and MVI performed data analysis; LGG-B, MD-R, HC-A, GG-O and JMB-M contributed all clinical data; MVI provided statistical supervision; LGG-B, CM-C and MAM-T provided research supervision; JLG-A, CM-C and MAM-T wrote the original draft; all authors provided expert input, reviewed and edited the final manuscript, and approved submission.

## ACKNOWLEDGEMENTS

JLG-A thanks an “Epriex” contract from “Plan Nacional de Recuperación, Transformación y Resiliencia” and UJI industrial fellowship. EC-M thanks the AECC fellowship. MAM.-T was supported by a ‘Maria Zambrano’ research contract (number MAZ/2021/03 UP2021-021) funded by the European Union-Next generation EU, UJI grant (23I469 UJI-2023-29) and Ramon y Cajal Research Fellow RYC2022-038481-by MCIN/AEI/ 10.13039/501100011033.

